# Probabilistic working memory representations in human cortex guide behavior

**DOI:** 10.1101/2025.11.17.688881

**Authors:** Ying Zhou, Clayton E. Curtis, Daryl Fougnie, Kartik K. Sreenivasan

## Abstract

Models of working memory make fundamentally different commitments to the architecture of individual memories. Information-sparse models conceptualize individual memories single point estimates agnostic to meta-cognitive variables such as uncertainty. In contrast, information-rich models propose memories re encoded s probability distributions over feature space that embed memory uncertainty in the shape of the distribution To distinguish these accounts, constructed probability distributions of memory from participant’s iterative reports of motion direction each trial. Remarkably, the idiosyncratic shape of these distributions (e.g., asymmetry) single trials matched the shape of neural probability distributions decoded from fMRI patterns measured from occipital and parietal cortex. Consistent with information-rich models, the neural representation of n individual memory encodes mor than the memorized feature; its variance (i.e., width and asymmetry) encodes idiosyncrasies whose read-out predicts memory behavior.

## Introduction

We know surprisingly little about the precise format by which working memory representations stored in the brain. For example, when temporarily maintain recently encountered direction of random dot motion in working memory, do w store probability distribution ver the space of all possible motion directions, or do we instead represent a specific direction of motion? The former view is inspired by probabilistic models that propose the activity of populations of encodes full probability distribution feature space^1–3^ We refer to these models *information rich* because the population of encodes memory content and memory uncertainty in multiplexed representational space akin to probability distribution ver possible feature values^4–6^ In contrast, high-threshold^7,8^ mixture^9,10^ and continuous attractor models^11,12^ ar *information sparse* because they represent memories s a single point estimate in feature space, with memory uncertainty potentially encoded in separate neural population. Information sparse representations preclude direct readout of uncertainty from the population storing the memory In standard attractor models of working memory^12^ for instance, memory is encoded by the increased response gain, *bump,* in units tuned to the memorized feature. While memory errors stem from the magnitude and direction of the bump’s drift ver time^11^ critically, n information about memory uncertainty can be discerned from the population activity. Ultimately, these two classes of models make fundamentally distinct claims about the format of memory representations, the mechanisms by which individual memories r transformed into behavioral output, and the origins of meta-cognitive information, such uncertainty. Thus, adjudicating between them is essential for understanding the neural basis of working memory.

Unfortunately, the ability to elucidate the architecture of individual memories is limited by experimental approaches that aggregate data over trials, which result in identical predictions for information-rich and -sparse models. Imagine scenario in which the same feature value is remembered several trials. Information-rich models would represent those memories full probability distributions with roughly the mean, but with distribution shapes that vary trials. Trial-wise idiosyncrasies in the shape of the distribution, such its asymmetry, may skew beliefs about the feature value in memory^13–15^ However, this information would be lost when averaging ver trials, resulting in roughly symmetric distribution centered on the memorized feature value (Figure 1A, left). Critically, the same distribution an result from aggregating point estimates generated by information-sparse models (Figure 1A, right), making it impossible to distinguish the two models. Thus, ignoring trial-wise idiosyncrasies obscure information-rich representations and prevent from fully understanding how the brain encodes memory (for related discussions, see^16,17^).

**Figure 1.**
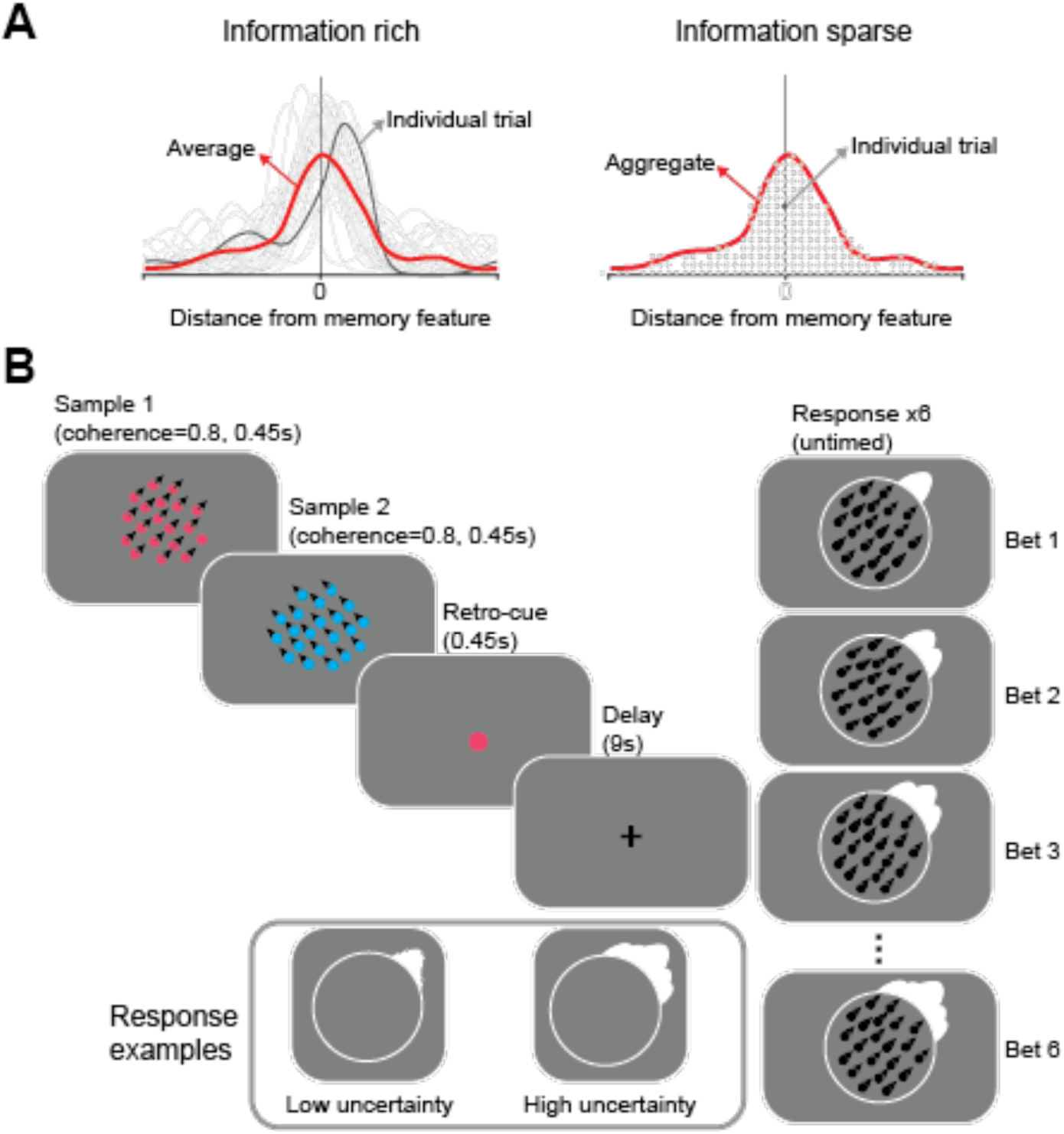
**(A)** Aggregated versus trial-wise representations Left: when individual memories are full probability distributions (each trial is indicated by gray line), predicted by information-rich models the trial-averaged distribution (indicated by the red line) is roughly symmetric and centered the memory feature. Right: when individual memories point estimates in feature space (each trial is indicated by gray dot), predicted by information-sparse models the trial-averaged distribution (indicated by the red line) is identical to the average distribution shown in the left panel Thus, information-rich and information-sparse models make indistinguishable predictions when data is aggregated trials **(B)** Trial schematic of the betting game task Participants presented with two dot-motion stimuli followed by retro-cue, and retained the cued motion direction memory delay period They reported the cued motion direction by placing six distributions around the probed circle to create an average distribution representing their memory on that trial The inset depicts example responses when participants had low high uncertainty about their memory. A video of the task is available at https://osf.io/zc3w4/

In the current study, w tested the hypothesis that ur memories are influenced by how uncertainty varies asymmetrically cros the feature space of n encoded memory This hypothesis predicts that the specific shape of neural probability distributions that encode individual working memories should predict idiosyncrasies of memory-guided behavior. To test this prediction, we derived estimates of neural probability distributions of individual memories using Bayesian decoder^18,19^ along with recently-developed procedure that allows participants to build probability distributions of their memory through sequential reports^13,14^ (Figure 1B). To preview, found that both behavioral and neural probability estimates highly idiosyncratic with asymmetric distributions feature space that diverged markedly from symmetric distributions derived from averaging trials. Critically, in several brain in occipital and parietal cortex, the shapes of the decoded neural distributions closely matched those of behavioral probability distributions on single trials. These results support information-rich models in which working memory content and uncertainty are encoded jointly within a single neural population as a complex probability distribution that sculpts memory-guided behavior.

## Results

### Decoding results

We collected fMRI data while participants performed working memory task in which they wer presented with two patches with moving dots and, after a memory delay, were asked to report the global motion direction of the cued patch (see Figure 1B and ***Experiment design*** for details). To identify individual memory representations encoded in the brain, utilized Bayesian decoder^20^ to derive posterior probability distribution the full motion space (0°–360°) from fMRI signals recorded during the delay period of individual trials (see Figure 2A and ***Bayesian decoding*** for details). Decoding wa conducted separately for regions of interest (ROIs; Figure 2B) in retinotopically organized regions in visual cortex (V1-V3, V3AB, hV4), lateral occipitotemporal (TO1/2, LO1/2), posterior parietal cortex (IPS0/1, IPS2/3), and lateral frontal cortex (sPCS).

**Figure 2.**
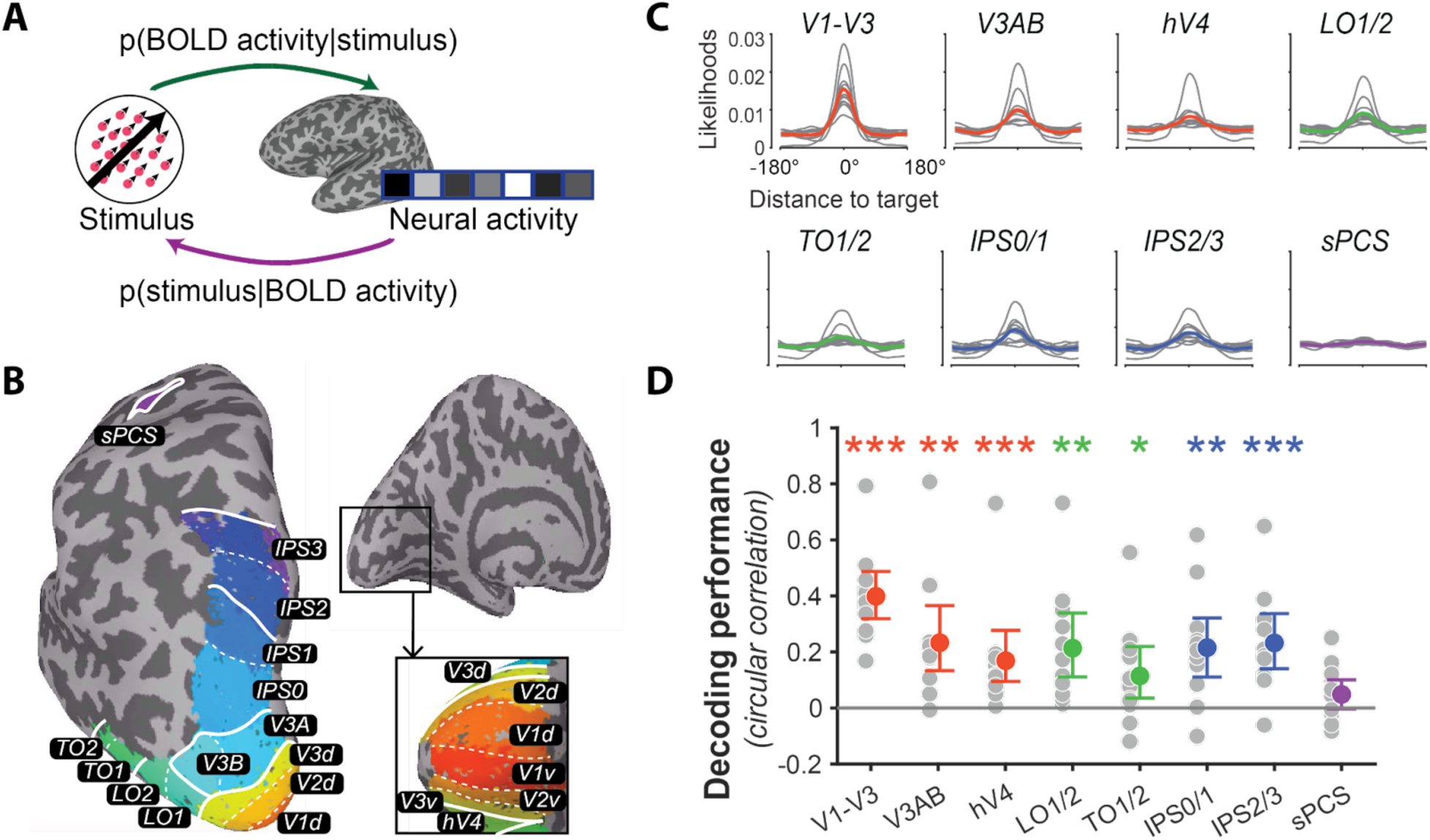
Decoding working memory representations. **(A)** Schematic of Bayesian decoding. TAFKAP first generates model to predict BOLD activity given the stimulus (p(BOLD activity|stimulus); green arrow) and then infers a full posterior distribution for the stimulus given the BOLD activity (p(stimulus|BOLD activity); purple arrow) **(B)** Topographically organized regions of interest (ROIs) shown for the left hemisphere of representative participant. Colors and labels indicate ROIs defined using probabilistic atlas of retinotopy ^27^ with boundaries between ROIs indicated by white lines ROIs sharing confluent fovea grouped together ^28,29^ **(C)** Decoded posterior distributions averaged over trials for each ROI Individual trial distributions aligned to reference of 0 and averaged trials Gray lines depict the centered posterior averaged all trials for individual participants; colored lines the group average Color indicates anatomical grouping of ROIs: red for early visual regions, green for lateral occipito-temporal cortex, blue for parietal cortex and purple for frontal cortex **(D)** Circular correlation between the decoded and target motion directions Gray dots represent individual participants and colored dots indicate the group for each ROI Stars indicate significant difference between the actual decoding performance and null simulation where the circular correlation calculated with target directions randomly permuted trials For all figures, bars indicate the 95% bootstrap confidence interval and stars denote significance (***: p .001; **: p .01; p .05, false discovery rate-corrected across ROIs with q=.05 where appropriate)

First, to compare findings to previous work, employed standard approach that aggregates decoding results cross trials. In each ROI, aligned the decoded distributions by setting the target to 0° and averaged the aligned distributions cros trials (Figure 2C). On each trial, w used the circular mean of the decoded probability distribution to represent the decoded motion direction^19,20^ Consistent with previous studies using similar decoding methods^19,21–26^ we able to decode memory content from occipital and parietal ROIs. To quantify decoding performance, calculated the circular correlation between the of the decoded distribution and target motion direction on individual trials. This correlation was significantly higher than null distribution generated by randomly permuting the target direction across trials. This finding consistent in all ROIs except sPCS (all correlations ≥ .11; *p* Figure 2D and Table S1). ≤ .018, Cohen’s *d* ≥ .66;

The shapes of averaged decoded posterior distributions were not, however, representative of single-trial decoded distributions. In particular, individual trial posteriors often exhibited marked asymmetry that was lost when the distributions were averaged across trials. As shown for an example participant and ROI in Figure 3, the shapes of individual trial posteriors were highly variable (Figure 3A) and exhibited a range of asymmetry that diverged from the trial-averaged level (Figure 3B). Of course, degree of trial-wise asymmetry is expected if the underlying neural representation is perfectly symmetrical due to noise inherent in the decoding process This noise is inherent because the voxel tuning used to compute likelihoods in the posterior distributions r estimates, not ground truth. To determine if the asymmetries we observed could be attributed to estimation noise, we compared the asymmetries in the decoded data to those obtained from simulations of randomly re-centered decoded distributions. This re-centering procedure mimics the outcome of decoding with completely uninformative tuning by shuffling the likelihoods for all possible values, thereby capturing the maximal asymmetry expected from estimation noise alone. We found that the observed asymmetries in the decoded distributions were significantly larger than the asymmetries in the simulations (*ps* ≤ .001, Cohen’s *d* ≥ 2.80; Figure 3C) confirming that they cannot be explained by estimation noise. Moreover, the observed asymmetries approached the theoretical upper bound estimated by randomly combining the left and right halves of posteriors from different trials, suggesting that the decoded representations r intrinsically and highly asymmetric (Figure 3C).

**Figure 3.**
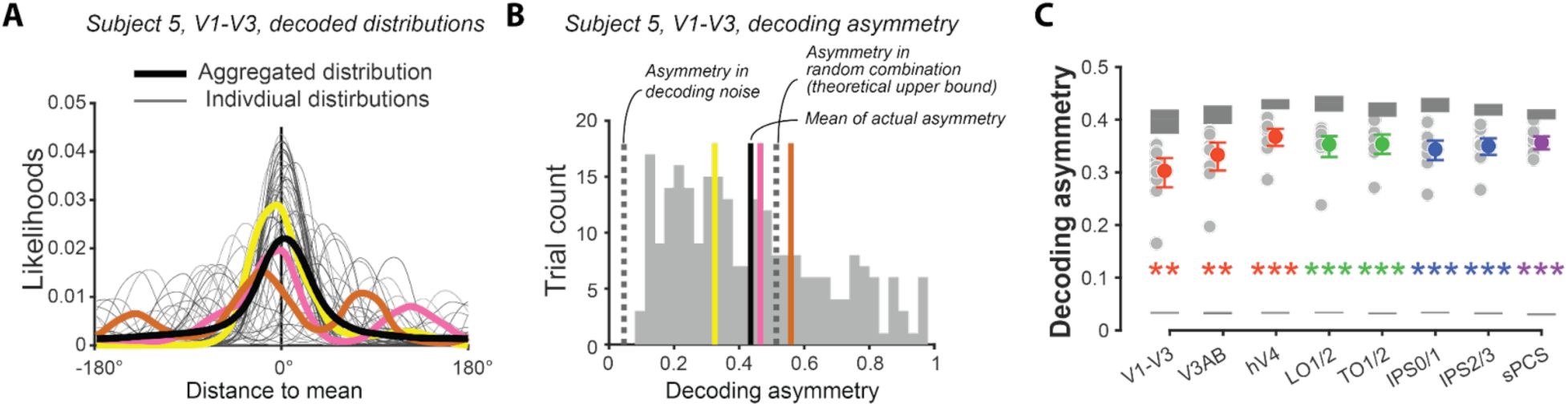
Asymmetry in decoded distributions **(A)** Averaged versus individual decoded distributions for sample participant and ROI The thick black line indicates the posterior distribution averaged trials, while the gray lines denote the decoded posterior distribution individual trials Posterior distributions for three example trials highlighted in yellow, pink, and brown **(B)** Asymmetry trials for the participant and ROI in A The light gray dashed line near 0 represents the asymmetry expected due to estimation noise alone (see **decoding performance** for details). The dark gray dashed line is the theoretical upper bound of asymmetry. The colored lines correspond to the example trials highlighted in panel A **(C)** Asymmetry ROIs. Gray dots represent individual participants and colored dots the of the measured asymmetry Light gray bars (bottom) indicate the amount of expected asymmetry from estimation noise, and dark gray bars (top) indicate the theoretical upper bound The heights of bars indicate 95% bootstrap confidence intervals Stars denote significant difference between measured asymmetry and asymmetry attributable to estimation noise.

Next, tested if the asymmetries in neural distributions meaningful by asking whether they predict asymmetries in memory-guided behavior. To do so, however, required behavioral task sensitive to the potential probabilistic nature of memory. Single reports about one’s memory are insufficient for this purpose. Thus, w employed our recently developed behavioral paradigm that use multiple reports about single memory item to estimate the distribution-level information contained in individual working memories^13,14^

### Betting game response distribution

On each trial, participants constructed distribution reflecting their memory by placing 6 bets about target motion direction instead of single report (see Figure 1B and ***Experiment design*** for details). To compare performance with traditional single-report tasks, we calculated the mean absolute error of the first response (Figure 4A). Error was lower for the first response (M 22.49°, SEM 2.70°) than subsequent responses (M 25.26°, SEM 2.60°; *p* .001, Cohen’s *d* 1.776), suggesting that the first response effortful best estimate of target identity. Moreover, the the first response comparable to that reported in prior single-report studies of working memory for motion direction^30^ suggesting that multiple responses did not lead to strategy shifts on the first response relative to single-response paradigms ^13^

**Figure 4.**
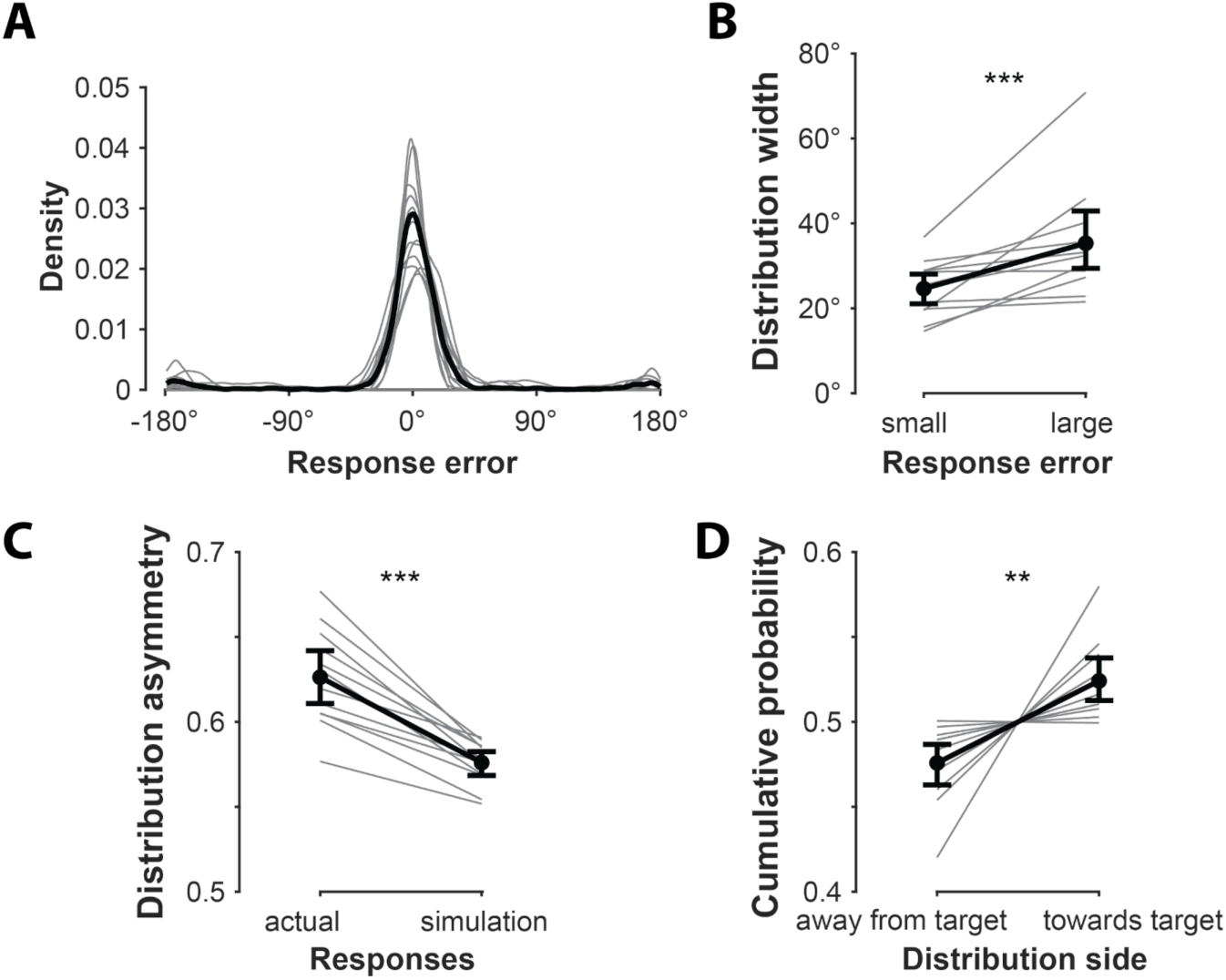
Behavioral performance. **(A)** The distribution of participants’ first response In this and following panels, gray lines indicate individual participants and the black line represents the average across participants **(B)** Distribution width in trials with small vs large the first response The bet distribution wider in trials with less accurate first responses consistent with the notion that participants indicating their memory uncertainty with the placement of their bets **(C)** Distribution asymmetry (the cumulative probability the larger side when the bet distribution centered relative to the first response). The actual behavioral asymmetry of bet distributions wa larger than that in simulated bet distributions where the signs of bets 2-6 relative to the first response randomly changed. **(D)** Target information in distribution asymmetry Cumulative probability greater toward the target side relative to the first response than the opposite side The stars in each figure indicate significant difference between the two conditions

A key question is whether bets 2-6 provide additional information about the target beyond the first bet, despite the fact that performance was individually worse for bets 2-6. For example, these extra bets may convey useful information about memory uncertainty; if participants’ memory about the motion direction is precise, they likely to cluster their bets, but if not, they likely to disperse their bets widely (see inset in Figure 1B). Consistent with this, found that trials with larger response errors were associated with broader bet dispersion (*p* .005, Cohen’s *d* = .83; Figure 4B; see also^19^).

Reminiscent of the neural decoded distributions, bet distributions were asymmetric around the first bet. That is, when centered on the first response, the cumulative probability was larger on side than the other. This asymmetry significantly larger than that expected due to random response noise, estimated by simulations in which the directions of subsequent responses wer randomly flipped relative to the first response on each trial (see ***Behavioral performance*** for details; M 0.63 vs 0.58; *p* < .001, Cohen’s *d* = 2.32; Figure 4C). Critically, this asymmetry contained meaningful information about the target: the cumulative probability on the target side (M 0.52) wa significantly larger than n the opposite side (M 0.48; *p* = .006, Cohen’s *d* = 1.07; Figure 4D). This asymmetry results in the of the bet distribution being faithful representation of the target direction (M 18.90°, SEM 1.77°) than the first bet (*p* .001, Cohen’s *d* = .999). These results demonstrate that the asymmetric shape of trial-wise response distributions, arising from participants’ subsequent bets, conveys information about individual working memory targets. Notably, this information would be lost and participants’ knowledge about their memories would be underestimated if only the first response wer collected, demonstrating the utility of multiple response paradigms.

### Correspondence between brain and behavior

In order to establish that the shape of our neural and behavioral distributions represents meaningful information about individual memories, it is essential to demonstrate correspondence between neural and behavioral estimates at the single-trial level We first replicated previous findings^19^ that the and width of neural estimates predict behavioral reports (Figure S1). However, this analysis ignores the full distribution shape, and therefore is insufficient to adjudicate between information-rich and -sparse models. To test correspondence between the shapes of our neural and behavioral estimates beyond the mean and width, we examined whether the neural distribution predicted the placement of bets individual trials better than the distribution reflected about its (Figure 5A). Importantly, flipping the neural estimate in this manner results in a distribution with the ame mean and width s the original, but with reversed asymmetry. Information-rich models predict that the original neural estimate would correspond with behavior better than the flipped estimate, while information-sparse models predict difference. Supporting information-rich models, found that the of the likelihoods the six behavioral responses in the neural distribution significantly larger than that in the distribution flipped around its in all ROIs except sPCS (*p*_s_ ≤ .010, Cohen’s *d*_s_ ≥ .829; Figure 5B, Figure S2). These results indicate that idiosyncrasies of probabilistic neural estimates, such as the asymmetry, in addition to their mean and width, can predict characteristics of memory-guided reports

**Figure 5.**
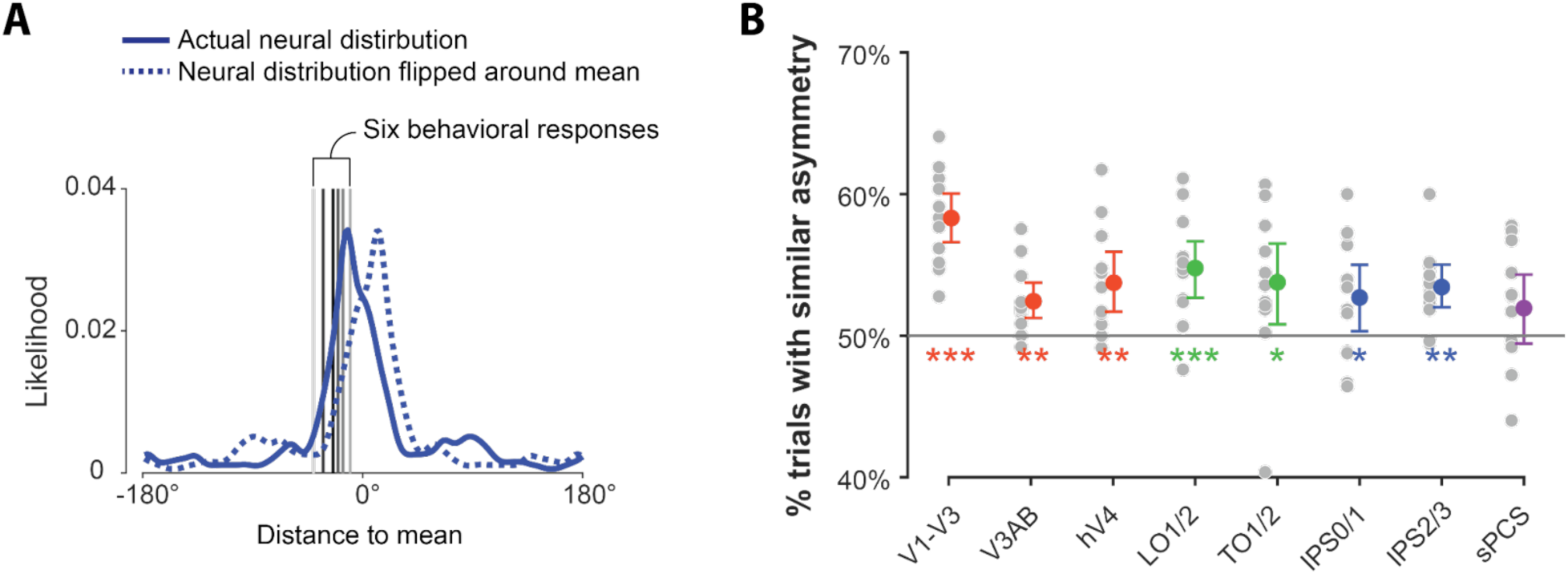
Brain-behavior correspondence. **(A)** Schematic of the asymmetry comparison between neural and behavioral estimates Filled blue lines represent the actual neural distribution centered at its for individual trial and dashed blue lines depict the neural distribution reflected about its Both distributions share the and width, differing only in their asymmetry Vertical gray lines from darker to lighter indicate behavioral responses 1-6 We compared the of likelihoods at the six behavioral responses for the actual and flipped neural distributions on individual trials **(B)** Brain-behavior correspondence was quantified as the percentage of trials which the actual neural distribution fit the behavioral response better than the flipped neural distribution Gray dots denote the correspondence for individual participants, and colored dots denote the of correspondence for all participants in different ROIs Stars indicate that the actual neural distribution fits the behavioral responses better than the flipped neural distribution

### Causes of asymmetries in individual memories

What asymmetries in memory? We evaluated two potential that predictable across trials, while acknowledging that other sources such as non-Gaussian noise^6,10^ may also contribute to memory asymmetries but r substantially more difficult to quantify. First, there is ample evidence that memories can be biased by information that was previously observed but is currently task-irrelevant. This includes uncued items from the current trial^31,32^ a well s previously relevant memory items from the preceding trial^33–35^ To ascertain whether task-irrelevant items contributed to the asymmetries observed, measured whether they systematically biased the neural and behavioral distributions In V3AB, the likelihood in the decoded probability distribution the uncued motion direction significantly smaller than the likelihood over the ame position in the distribution flipped around the mean (*p* = .004, Cohen’s *d* = .860; Figure 6A, left). There was corresponding reduction in the size of the behavioral distribution toward, compared to away from, the uncued motion relative to the first bet (*p* = .023, Cohen’s *d* = .674; Figure 6A, right). Together, these results consistent with the idea that inter-item competition from uncued memory items creates repulsive bias in memory representations^36^ and suggest that this bias contributed to observed asymmetries in neural and behavioral probability distributions. When we repeated the same analysis for the cued motion direction from the previous trial, we observed similar repulsive bias in the decoded distribution in V1-V3; however, this effect did not survive multiple comparisons (*p* = .064, Cohen’s *d* .803). No bias wa observed for the other ROIs (*ps* ≥ .107, Cohen’s *ds* ≤ .360) or the behavioral (*p* .200, Cohen’s *d* = .237) distributions (Figure 6B), perhaps due to the long interval between trials^37,38^

**Figure 6.**
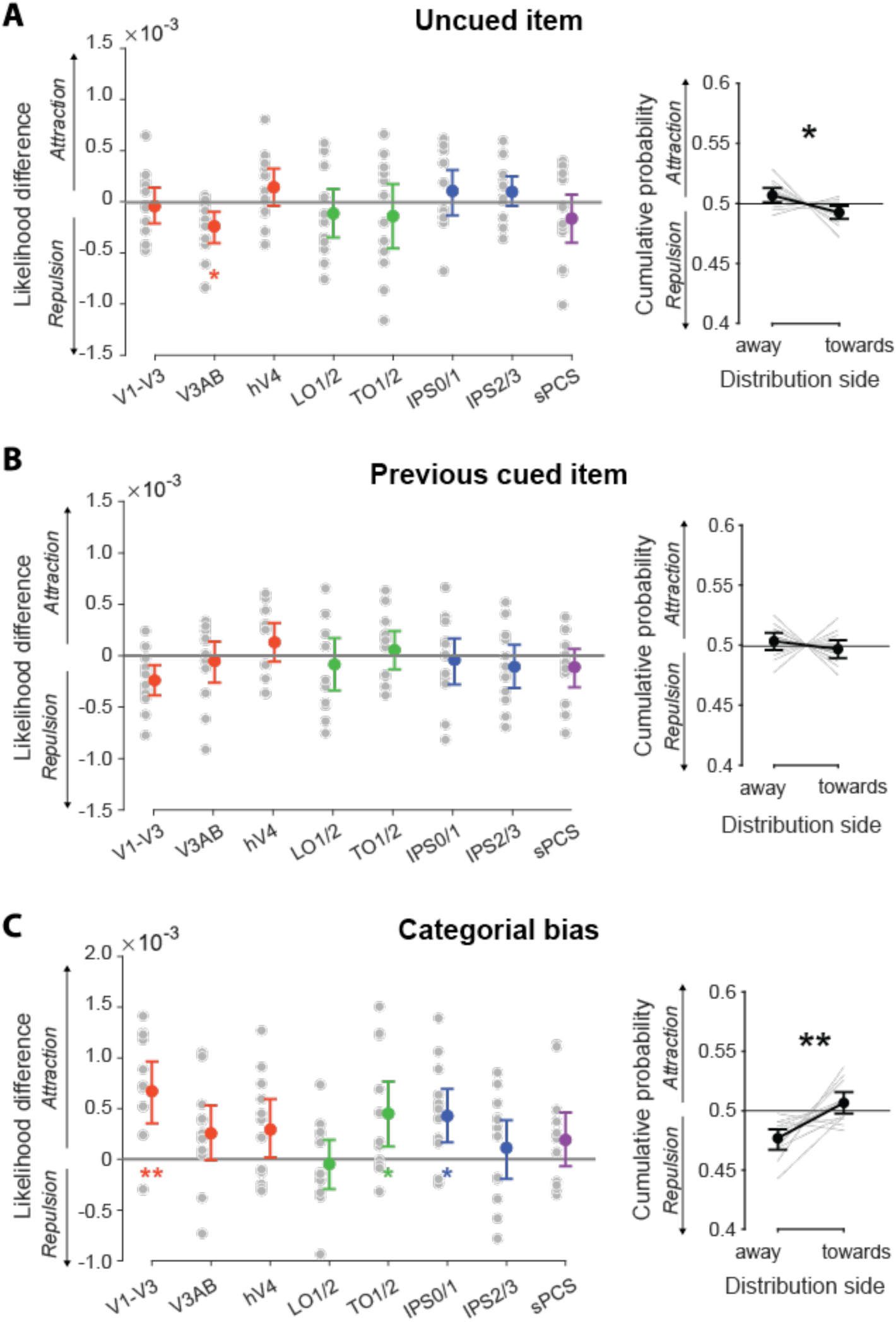
Causes of asymmetries in neural and behavioral distributions **Left column** Neural bias quantified by calculating the likelihood the motion direction of interest and subtracting the likelihood over the same position in the neural distribution reflected about its Positive (negative) values indicate attractive (repulsive) bias, respectively Gray dots represent individual participants, colored dots indicate the group for each ROI, and stars indicate bias that differed significantly from **Right column** Behavioral bias, quantified the difference between the cumulative probability the side of the distribution away from toward the motion direction of the interested cause relative to the first response Gray lines represent individual participants, the group is shown in black and stars indicate significant difference between the distribution sizes the two sides **(A)** Bias caused by the uncued motion direction the trial Left: There significant repulsive bias away from the uncued direction in V3AB Right: There significant repulsive bias from the uncued direction in the behavioral distributions. **(B)** Bias caused by the cued direction the previous trial Left: A trend towards repulsive bias in V1-V3 did not survive FDR correction Right: Despite a trend towards a repulsive bias (7 of 12 participants) no significant bias was observed in the behavioral distributions **(C)** Categorial bias caused by nearby canonical directions Left: There significant repulsive bias towards the nearby canonical directions in V1-V3, TO1/2, and IPS0/1 Right: There significant attractive bias toward the nearby canonical directions in the behavioral distributions

Another factor that could contribute to asymmetries in memory is systematic categorical bias toward canonical directions (vertical, horizontal, and oblique)^39,40^ Supporting this idea, w found that decoded probability distributions in V1-V3 (*p* .001, Cohen’s *d* 1.170), TO1/2 (*p* .015, Cohen’s *d* .752), and IPS0/1 (*p* .006, Cohen’s *d* .807) had a greater likelihood ver nearby canonical directions than the opposite side relative to the The bet distribution similarly larger the side facing the nearby canonical directions than the opposite side relative to the first bet (*p* < .001, Cohen’s *d* 1.080; Figure 6C and Figure S3). Thus, attraction to canonical feature values may also contribute to memory asymmetry. The asymmetry w observed is not just limited to the elements tested here, but is likely the result of complex interplay of factors, such as trial-wise variability^41^

## Discussion

Here, tested the hypothesis that asymmetrical uncertainty mnemonic feature space shapes neural probability distributions, which in turn predicts idiosyncrasies in memory-guided behavior. Our findings support this hypothesis and demonstrate that memory and uncertainty re jointly represented cros the visual hierarchy and that the architecture of individual memories is complex, probabilistic in nature, and drives memory-guided behavior. These results represent mutual validation of single-trial behavioral and neural probabilistic memory estimates, and support information-rich models of working memory positing that multiplexed information about memory and uncertainty is encoded in the ame neural populations. One way to store multiplexed information in neural populations is proposed by the theory of probabilistic population codes: the brain knows how patterns of neural activity relate to visual features and uses population activity to estimate both the likely feature and the uncertainty that comes with it^1,3,42–44^ In contrast, information-sparse theories conceive of memories point estimates in feature space that lack notions of uncertainty. Some information-sparse models, such continuous attractors^11,12,45,46^ can in principle incorporate information about memory uncertainty; in such cases, our findings place major constraints on precisely how uncertainty is represented. For example, models where uncertainty is encoded separately from memory content but influences the representation through feedback mechanisms ^47,48^ are inconsistent with our observations any information about asymmetry would be lost and could not influence behavior. Put another way, asymmetric uncertainty suggests that memory and uncertainty jointly encoded.

Furthermore, our framework of trial-wise probabilistic brain-behavior comparison provides a critical tool for differentiating the contributions of distinct brain regions to working memory. While memory features can be simultaneously decoded from multiple brain regions including visual, parietal, and prefrontal cortices^19,49–51^ (see^52,53^ for reviews), precise understanding of each region’s role remains elusive^54–56^ Importantly, it is misleading to infer that these regions encode information of the quality based single estimates (e.g., width) of memory; even if the neural probability distributions estimated from these rea have identical mean and width, differences in their shape could signify meaningful differences in the information that they represent. Our approach, which identifies nuanced but behaviorally relevant aspects of the shape of individual memory representations, offers a powerful means for distinguishing the quality of working memory representations in different brain regions, and is thereby able to clarify their unique functional roles. Our results suggest that early visual areas, often considered less important for working memory^55,57,58^ contain sufficient information to guide complex memory-guided behaviors involving uncertainty and bias.

The field’s reliance on simple reports and aggregate analyses is partly based on the assumption that such methods adequately characterize useful working memory properties. For example, average response rror cross set sizes has been used to infer working memory capacity limits^9,59,60^ However, there several problems with this perspective. First, results argue that the enterprise of characterizing average working memory state may miss the point; memory states highly variable trials, with few resembling the aggregate (Figure 3A). Rather than illuminating purported qualitative average memory state, aggregate approaches may mask the underlying mechanisms that produce working memory representations^61^ Second, there is no established theory explaining how uncertainty in neural memory representations is converted into behavioral reports, despite being an essential inferential step. One might assum that participants take optimal sample from memory, such the of memory distribution. However, found that averaging multiple behavioral reports accurate than individual reports, suggesting that each individual sample is suboptimal. Behavioral responses may instead reflect noisy or partial samples from the underlying probabilistic representation^13,62,63^ Alternatively, the underlying distributions may be too complex for one report to summarize^6^ These results underscore the need for theory of how neural uncertainty in memory is converted into behavioral reports (an typically relegated to decision-making^64^), in order to make meaningful advances in linking brain and behavior.

Our work highlights several key considerations for future research. First, examining working memory representation on a per-trial basis is essential, as this approach can reveal additional information, such trial-specific uncertainty, that is lost in aggregated analyses. Second, individual trials, relying single point estimate either summary of neural decoding summary of behavioral report is insufficient; instead, estimating the full probability distribution can provide a more comprehensive understanding of working memory representation and memory-guided behavior. Finally, tracking brain-behavior relationships at the single-trial level is crucial for deepening our understanding of working memory^16,65,66^ By adhering to these principles, we constrain working memory models by highlighting that successful theories must account for the joint encoding of both mnemonic and metacognitive information. Further, show that deviations from the aggregate have consequences they make successful predictions about what participants will report and with how much certainty they report it.

## Methods

### Participants

All procedures were approved by the New York University Abu Dhabi (NYUAD) Institutional Review Board. Based on sample sizes from previous fMRI studies using Bayesian decoding^18,19,30,67^ recruited fifteen neurologically healthy volunteers (5 males, 27.5 ± 4.5 years old) from the university community. All reported normal corrected-to-normal visual acuity and normal color vision. Two participants excluded from subsequent analysis because their response error exceeded two standard deviations above the group mean, and on withdrew from the study before completing data collection. The remaining 12 participants (5 males, 28.7 ± 4.1 years old) completed two (2 participants) or three (10 participants) MRI sessions. Participants compensated for their time with AED 120 (∼USD 33) per session, plus a performance-based bonus of up to AED 50 (∼USD 13).

### Experimental design

We generated stimuli and interfaced with the MRI scanner, button box, and response dial using MATLAB software (The MathWorks, Inc., Natick, MA) and Psychophysics Toolbox 3 (Brainard, 1997). Stimuli were presented using PROPixx DLP LED projector (VPixx Technologies, Inc., Saint-Bruno, QC, Canada) located outside the scanner room and projected through waveguide and onto translucent located at the head of the bore. Participants viewed the at viewing distance of 88 through mirror mounted the head coil. A scanner trigger synchronized stimulus presentation and image acquisition.

Each trial (Figure 1B) began with two sequential colored motion patterns (80% coherence; radius 4 degrees of visual angle [dva]) presented for 450 ms each, separated by 200 m interval. Each motion pattern comprised 70 moving dots (0.16 dva) traveling at 9 dva/s, with motion directions drawn, without replacement, from nine evenly spaced categories (20°–360°), jittered ±5° to prevent participants from forming categorical representations of the motion. Participants memorized both motion directions for later recall. After 200 ms, 450ms color retrocue (0.2 dva) indicated which motion direction to retain. Following 9000 ms delay, participants adjusted a black probe motion pattern (100% coherence) using an MR-compatible dial (Current Design, Inc., Philadelphia, PA) to match the cued direction and confirmed their response with a button press

Instead of single response, in typical continuous report working memory tasks, participants made 6 bets to report the target motion direction each trial^13,14^ Each bet displayed von Mises distribution (circular SD 10°) that moved on the creen a participants adjusted the dial, and the sum of the six bets formed the final response distribution for that trial. Participants could spread the bets freely, with the cumulative distribution updated in real time to reflect all bets placed o far plus the new bet reflected by the current dial position. A video of the task is available at https://osf.io/zc3w4/

To encourage accurate reporting of something resembling internal uncertainty, points awarded based on the height of the final response distribution over the target motion direction:

Points = Height of final distribution at target direction * 500.

The first bet was assigned double the height of bets 2-6 (i.e., worth twice as many points) to encourage accuracy n the first bet. To discourage participants from only stacking bets, w implemented diminishing rewards for stacking. Specifically, if the center of the bet is at *b* the existing cumulative response distribution is *y* and is the Mises distribution representing the new bet, then the new cumulative response distribution *y*’ after the new bet is: 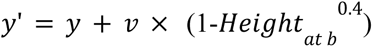, where 0.4 is the penalty parameter.

Thus, the higher the existing cumulative distribution at the new bet’s location, the smaller the incremental gain in height after adding that bet. This penalty was visually represented in real time within the response display. Consequently, to maximize their points (and thus bonus payment), participants incentivized to cluster bets when confident about the target motion direction and to spread them when uncertain (see Figure 1B, inset).

After participants placed all six bets, 1500 m feedback screen displayed the final response distribution, the target direction, the points earned on that current trial, and the cumulative total. The inter-trial interval was 10500 ms. Each scanning session consisted of 9 or 10 runs of 9 trials.

### fMRI Methods

*MRI data acquisition* Scanning sessions conducted at the NYUAD Brain Imaging Core Siemens Prisma (3T) MRI canner with 64-channel head/neck coil. Functional BOLD can were acquired using n EPI pulse sequence with 44 slices and a voxel size of 2.5 isotropic (4x simultaneous-multi-slice acceleration; FoV 200 200 mm, n in-plane acceleration, TE/TR 30/750 ms, flip angle = 50°, bandwidth = 2604 Hz/pixel, 0.51 ms echo spacing, P A phase encoding). To correct for local spatial distortions, estimated field map of the field inhomogeneities by acquiring pairs of spin echo images with normal and reversed phase-encoding directions with identical slice prescription to the functional data and simultaneous-multi-slice acceleration (TE/TR = 36/3100ms, 3 volumes per phase encoding direction). To enable precise localization of functional data, we collected T1-weighted whole-brain anatomical cans using MP-RAGE sequence with 208 slices and voxel size of 0.8 isotropic (FoV 256 240 mm, TE/TR = 3.45/2400 ms, flip angle = 8°, bandwidth 220 Hz/Pixel). Slice positioning and FoV individually adjusted to full-brain coverage. To further improve the surface extraction, acquired additional 3D T2-weighted SPACE sequences in the same resolution as the T1-weighted anatomical can (FoV 256 240 mm, TE/TR = 564/3200 ms, bandwidth = 744 Hz/Pixel).

*MRI data preprocessing* Our preprocessing followed the <u>Minimal Preprocessing Pipeline of the</u> <u>Human Connectome Project</u> (MPP)<u>(Glasser et al. 2013)</u> which optimized for NYUAD’s High Performance Computing cluster to multiple participants in parallel. Briefly, the structural MRI (T1 and T2) aligned, brain extracted, and corrected for readout distortions before being projected into MNI space using Boundary Based Cross Modal Registration (BBR), and then finally bias-field corrected using the combination of T1 and T2 contrasts. Functional data was corrected for using MPP’s *fMRIVolume* pipeline. After correcting scanner-specific gradient distortions, the BOLD time series underwent motion correction through rigid body registration to align with the single-band reference gradient echo EPI volume. Susceptibility artifacts corrected utilizing the pair of Spin Echo EPI volumes with opposite phase encodings (AP and PA directions). Subsequently, BBR was used to co-register the single-band reference gradient echo EPI volume with the corrected T1w volume. This registration transformation was then consolidated into a single transformation step that was applied to the BOLD series. Since the volumetric processing of the MPP pipeline only produces corrected functional in MNI space, used FSL’s *applywarp* algorithm to resample the BOLD data into native space. We visually inspected the overlap of the native BOLD data with the native corrected T1w and T2w volumes to ensure the accuracy of the registration. All analyses were conducted in corrected native volume space. Finally, we removed linear trends from the BOLD series and normalized (z-score) across all time points within each run.

*Regions of interest (ROIs)* We used FreeSurfer (http://surfer.nmr.mgh.harvard.edu/) to construct brain surfaces from the corrected T1w anatomical using the *recon-all* command, and identified the following participant-specific ROIs in surface space from the probabilistic map of visual topography^27^ using the *atlas* command in Neuropythy^68^ hV4, V1, V2, V3, V3A, V3B in occipital visual cortex; LO1, LO2, TO1, TO2 in lateral occipito-temporal cortex; IPS0, IPS1, IPS2, IPS3 in parietal cortex; and sPCS in frontal cortex. We combined ROIs that share the same foveal confluence^28,29^ (V1, V2, and V3; V3A and V3B; IPS0 and IPS1; IPS2 and IPS3; LO1 and LO2; TO1 and TO2; Figure 2B). Finally, these ROIs were projected into native volume space.

### Bayesian decoding

In order to decode the motion direction stored in working memory n each trial, w used the averaged z-scored BOLD response for each voxel from 6.75s to 9s after the onset of the delay period the input to a Bayesian decoder, TAFKAP^20^ The output of TAFKAP is a posterior probability distribution of each stimulus value the whole feature space. Specifically, it first testing data to create generative model to predict the neural activity given stimulus value, then infers the posterior of stimulus value given the neural activity by applying Bayes’ theorem on unlabeled testing data (see the illustration in Figure 2A).

In the generative model, the voxel response given the stimulus (i.e., the target motion direction) was modeled as a multivariate normal distribution. Specifically, the average response of each voxel 𝑏_𝑖_ was determined by its tuning function 𝑓_𝑖_(*s*) plus a random noise ε_𝑖_

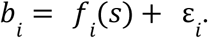

The tuning function 𝑓_𝑖_(*s*) of each voxel 𝑖 approximated by weighted of nine basis functions 𝑔_𝑘_() evenly distributed across the feature space:

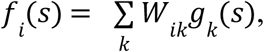

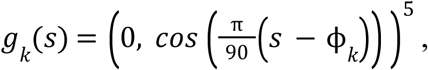

where 𝑔_𝑘_(*s*) is the 𝑘-th basis function, which peaks at direction ϕ_𝑘_and 𝑊_𝑖𝑘_ is weighting matrix that determines the weights of each basis function 𝑔_𝑘_(*s*) for each voxel 𝑖 The noise _𝑖_ for each voxel is assumed to be correlated between voxels, with covariance Ω such that ε ∼ 𝑁(0 Ω) and 𝑏_𝑖_𝑁(𝑓_𝑖_(*s*) Ω) The probability of multivoxel activity pattern 𝑏 = [𝑏_𝑖_]^𝑇^ is therefore given by:

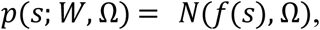

where 𝑓(*s*) [𝑓_𝑖_(*s*)] are the values of each voxel’s tuning functions given stimulus and voxel weight 𝑊 and covariance structure Ω are the free parameters needed to be estimated.

The free parameters were estimated from the data in leave-one-run-out cross-validation procedure. That is, each participant’s dataset divided into training and testing samples, such that each served the testing sample The training samples used to estimate the free parameters in the generative model. Specifically, the voxel weight 𝑊 was estimated by

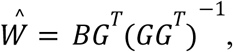

Where 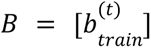 is matrix of multivoxel activity in all training samples and 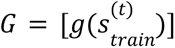is matrix of the values for the nine basis functions given stimulus in the training samples, and indexes trials. The covariance matrix Ω be theoretically estimated by a sum of noise independent between voxels and noise shared among all voxels,

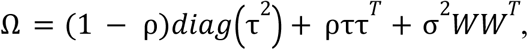

where 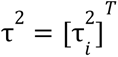 are the variance parameters of noise that is independent between voxels, and ρ is global correlation parameter for noise that is shared among all voxels; σ^2^ is variance parameter for noise that is shared between voxels with similar tuning properties; and 𝑊𝑊^𝑇^ indicates the similarity between tuning In addition to this theoretical covariance matrix, the model also considered the empirical sample variance, which can be calculated by

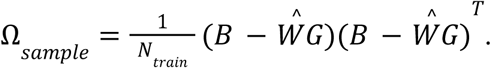

Considering both the theoretical and empirical covariance, the covariance matrix Ω_𝑛𝑒𝑤_ was modeled the sample covariance matrix “shrunk” to the theoretical covariance matrix Ω degree of shrinkage is determined by a free parameter λ

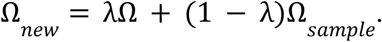

The parameters of this covariance model (ρ 𝑎𝑛𝑑 λ) were estimated by numerically maximizing their likelihood under the training samples, conditioned on the estimate of 𝑊 After estimating the free parameters using the training samples, inferred the posterior of stimulus value given the multivoxel responses in each testing trial by applying the Bayes’ theorem:

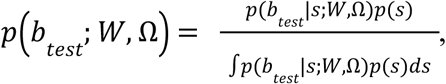

where a flat prior 𝑝() reflecting the equal possibility of the nine stimulus categories from which the target values sampled in the experiment, and the normalization constant in the denominator computed numerically the whole feature space We used discrete approximation of the posterior, evaluating 𝑝_(_𝑏_𝑡𝑒𝑠𝑡_ 𝑊 Ω_)_ at 180 equally spaced directions in the whole feature space (0°–360°).

To account for the uncertainty of model parameters, TAFKAP bootstrap aggregating method. We resampled the trials in the training data with replacement multiple times to generate many resampled training datasets. Each resampled training dataset had the same number of trials the training samples. For each resampled training dataset *j* set of free parameters 𝑊_𝑗_ 𝑎𝑛𝑑 Ω_𝑗_ estimated using the method mentioned above. The posterior distribution of stimulus given the multivoxel response in each testing trial in iteration *j* of bootstrapping was then computed as

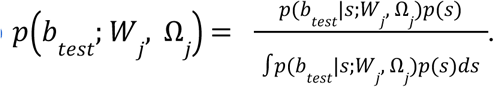

For each testing trial, this decoding performed multiple times based the parameters estimated using each resampled training dataset. The final decoding result of each testing trial was the averaged posterior distributions across all the bootstrapping iterations

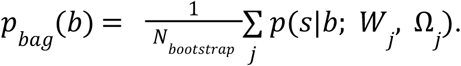

The number of bootstrapping iteration (𝑁_𝑏𝑜𝑜𝑡𝑠𝑡𝑟𝑎𝑝_) was determined by a stopping criterion based Jensen-Shannon divergence. In testing process, if the largest Jensen-Shannon Divergence between the new and the last 100 posterior distribution samples was smaller than 1e-8, or the bootstrapping iterations were larger than 5 10^4^, we terminated the bootstrapping process.

We smoothed the final posterior distribution with sliding window of 10° to facilitate comparisons with the bet distributions, which summations of Mises distributions with circular standard deviation of 10°.

### Decoding performance

For each trial, defined the decoded motion direction the circular of the posterior distribution obtained from the Bayesian decoder^19,20^ We assessed decoding performance using the circular correlation between the decoded and the actual motion directions^19,20^ For each participant and ROI, we computed the circular correlation and compared it to null distribution obtained by randomly permuting the target direction over trials and then recomputing the circular correlation 1000 times. We report the *p*-values at the individual participant level a the proportion of null circular correlations larger than the actual circular correlation in Table S1. For group-level inference for the circular correlation results, used nonparametric permutation-based test^26^ because there *priori* to believe that the data would be normally distributed. We compared the *t-*statistic derived from the group *t*-test of the actual circular correlation against 0 to null distribution of *t-*statistics obtained by comparing the null correlations for all participants in each permutation with 0 The *p-*value on group level wa the proportion of null *t*-statistics larger than the actual *t*-statistic, which are reported in Figure 2D. The effect size is reported as Cohen’s *d*^69^

The asymmetry of the decoded posterior distribution around its quantified by the dissimilarity caused by flipping the distribution If distribution is asymmetric around its mean, flipping it around the mean would result in a distribution dissimilar to the original distribution; in contrast, flipping a symmetric distribution around the mean would result in an identical distribution. We computed the overlap between the original and flipped distributions to measure the similarity between them. This has the advantage of not requiring any distributional assumptions such symmetry, unimodality, and well-established parametric forms, and has been shown to be able to considerably improve the interpretability of data analysis results in psychological research^70^ The overlap η between distribution 𝑓*_A_*(*x*) and distribution 𝑓*_B_*(*x*) was determined by

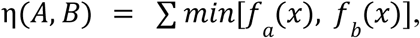

which is larger for more similar distributions^71^

To confirm that the asymmetry in decoded posterior distributions not simply caused by the noise in estimating voxels’ tuning curves, compared the actual asymmetry with simulations where voxels’ turning curves were random noise without any meaningful information. For each participant and ROI, the simulation was generated by re-centering the posterior distributions at randomly permuted locations and taking the mean of all these re-centered distributions. The re-centered distributions used to quantify the asymmetry of the simulated distribution described above for the actual posterior distributions. This process repeated 1000 times to derive null distribution of asymmetry. To compare the actual asymmetry with that caused by estimation noise and get the statistical results reported in Figure 3C, w used the nonparametric permutation-based test described above to compare the actual *t*-statistic derived from comparing the actual asymmetry for all participants with 0 to the null distribution of *t-*statistics obtained by comparing the asymmetry of estimation noise for all participants with 0 in each permutation. The *p*-value at the group level the proportion of null *t*-statistics larger than the actual *t*-statistic.

To further contextualize our asymmetry measure, we compared the actual asymmetry with the theoretical upper bound of asymmetry, which was simulated by randomly combining the left side of the distribution on each individual trial with the right side of the distribution on another trial when all distributions were centered at their mean, and recomputing the dissimilarity caused by flipping. This process repeated 1000 times for each individual trial and the average taken the theoretical upper bound asymmetry for each individual trial. The theoretical upper bound asymmetry for each participant and ROI was the average of all individual trials.

### Behavioral performance

We quantified behavioral response error as the circular distance between the first response and the target stimulus in previous studies using single response paradigms (Figure 4A). To demonstrate that the subsequent responses individual trials incorporates information about memory uncertainty, compared the width of bet distribution (the absolute value of the difference between the minimum and maximum bet error on a given trial) in accurate and inaccurate trials (Figure 4B). Trials were labeled accurate or inaccurate for each participant based on a median split of the error of the first response The group-level statistical comparison of bet distribution width between accurate and inaccurate trials calculated using nonparametric permutation-based test. We first performed paired *t*-test participants between accurate and inaccurate trials. The resulting *t*-statistic compared against null distribution generated by randomly permuting the trial labels (accurate vs inaccurate) within each participant and recomputing the *-*statistics 1000 times. The group level *p*-value was the proportion of null *t*-statistics larger than the observed *t*-statistic.

To quantify the asymmetry of the bet distribution, w used the cumulative probability on the larger side of the bet distribution after centering it the first response (Figure 4C). To confirm that the asymmetry created by subsequent responses not simply artifact of response noise (i.e., that randomly placed later bets could, by chance, produce apparent asymmetry around the first response), we compared it with a simulation where the direction of subsequent responses relative to the first response n individual trials wa randomly switched. The simulation process was repeated 1000 times for each participant to yield null distribution of asymmetry levels. We report group-level statistics from the nonparametric permutation-based test comparing the actual and simulated asymmetry levels, as described above.

To confirm the asymmetry of the bet distribution contained meaningful information about the target direction, compared the cumulative probability the side containing the target with that on the other side when the bet distribution was centered relative to the first response (Figure 4D). Group-level statistics were computed using a nonparametric permutation-based test comparing the cumulative probability on the target and non-target sides, as described above.

### Brain-behavior correspondence

We first tested brain-behavior correspondence in the and width of the distribution estimates by computing binned-correlation between the mean and width of behavioral and neural estimates a in^19^ For each participant, trials were sorted into four bins according to decoding rror The behavioral response rror wa computed by averaging cros trials within each bin We then pooled data points across participants (four data points per participant ne per bin) after removing the of each participant. Pearson correlation between the neural and behavioral estimates then computed based the pooled data (Figure S1). We compared the correlation coefficients to the null distribution obtained by permuting the data points in the pooled dataset to obtain a *p*-value. For visualization, we added the mean of each participant back to the data when plotting the binned-correlations. We repeated the same binning analysis using uncertainty instead of error.

To confirm the brain-behavior correspondence in the shape of the distribution estimates, examined whether the asymmetry of the neural decoded distribution could predict behavioral responses Specifically, tested whether the actual neural decoded distribution fits the behavioral responses better than the same neural distribution flipped around its mean For each individual trial, the likelihood of getting the actual 6 behavioral responses based n the neural distribution was calculated by

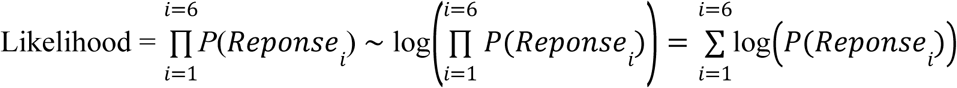

where P is the likelihood of 𝑟𝑒𝑝𝑜𝑛𝑠𝑒_𝑖_ at the neural distribution. If the likelihood of the actual decoded distribution is higher than that of the flipped decoded distribution in individual trial, this trial was labeled as a matching trial. The brain-behavior correspondence for each ROI and individual participant was calculated by the percentage of matching trials.

To generate chance-level of brain-behavior correspondence, we shuffled the distribution labels (actual flipped) and recomputed the brain-behavior correspondence for each participant and ROI. This process repeated for 1000 times to generate null distribution. To compare the actual brain-behavior correspondence with the chance-level and obtain the statistical results reported in Figure 5B, we used the nonparametric permutation-based test described above to compare the actual *t*-statistic derived from comparing the actual brain-behavior correspondence for all participants with 0 to the null distribution of *t-*statistics obtained by comparing the chance-level brain-behavior correspondence for all participants with 0 in each permutation. The *p*-value at the group level was the proportion of null *t*-statistics larger than the actual *t*-statistic.

To rule out the possibility that the brain-behavior correspondence in asymmetry was driven by trials where the mean of the neural distribution was poor predictor of behavior, we tested whether the relationship between neural and behavioral asymmetry held when the mean of the decoded distribution closely matched that of the behavioral distribution. For each ROI and participant, selected trials where the absolute distance between the neural and behavioral within standard deviation of the distance all trials (see Figure S2A for the percentage of selected trials). We then repeated the brain-behavior correspondence analysis described above on these trials (Figure S2B).

### Causes of asymmetries in individual memories

To investigate the of asymmetries in memory, examined the influence of task-irrelevant items and canonical motion directions (vertical, horizontal, oblique) decoded posteriors and the distribution of bets. The task irrelevant items we focused on were the uncued motion direction on the same trial and the cued motion direction from the previous trial. Canonical directions were defined as 90°, 135°, 180°, 225°, and 270°, and our analysis focused on the influence of these directions on targets belonging to the nearby categories of 100°, 140°, 180°, 220°, and 260°, respectively.

Neural bias quantified by finding the likelihood the motion direction of interest in the decoded probability distribution and subtracting the likelihood the position in the flipped distribution (i.e., the distribution reflected over its mean). To establish a chance-level of bias we shuffled the distribution labels (actual vs flipped) and recomputed the bias, iterating this process 1000 times to generate a null distribution for each participant and ROI. To evaluate the statistical significance of bias and obtain the results reported in Figure 6 and Figure S3, applied nonparametric permutation-based test previously described, comparing the actual *t*-statistic from the actual bias for all participants against 0, with the null distribution of *t*-statistics derived from comparing the chance-level bias for all participants with 0 across each permutation. The *p*-value at the group level wa calculated a the proportion of null *t*-statistics exceeding the actual *t*-statistic.

For behavioral bias, we first centered the distribution at the first bet, as was done for the asymmetry analysis of bet distributions in ***Behavioral performance*** Then compared the cumulative probability the side containing the motion direction of interest with that the opposite side. We report group-level statistics from the non-parametric paired *t*-test comparing the cumulative probability on the two sides as described above in ***Behavioral performance***

## Data and Code Availability

Data, experiment and analysis code, and other research materials associated with this study publicly available the Open Science Framework (OSF) project page https://osf.io/zc3w4/ The data comprises deidentified processed fMRI (timeseries from each voxel of each ROI).

## Author Contributions

Y.Z., D.F., and K.K.S. designed the experiment and study. Y.Z., C.E.C., D.F., and K.K.S. implemented the study. Y.Z. collected the data. Y.Z. analyzed the data with input from D.F. and K.K.S. Y.Z., C.E.C., D.F., and K.K.S. drafted and revised the paper. All authors provided critical feedback and approved the final paper for submission.

## Acknowledgements

This work wa supported by the NYUAD Center for Brain and Health, funded by Tamkeen under NYUAD Research Institute grant CG012 (K.K.S.) and NIH R01 EY-016407 and R01 EY-033925 (C.E.C). This research was carried out on the High Performance Computing resource at NYUAD. We thank Haidee Patterson for help with MRI data collection. We also thank Jorge Naranjo and Dr. Osama Abdullah for adapting the HCP preprocessing pipeline for use on NYUAD’s high-performance computing cluster.

## Extended Data

**Figure S1.**
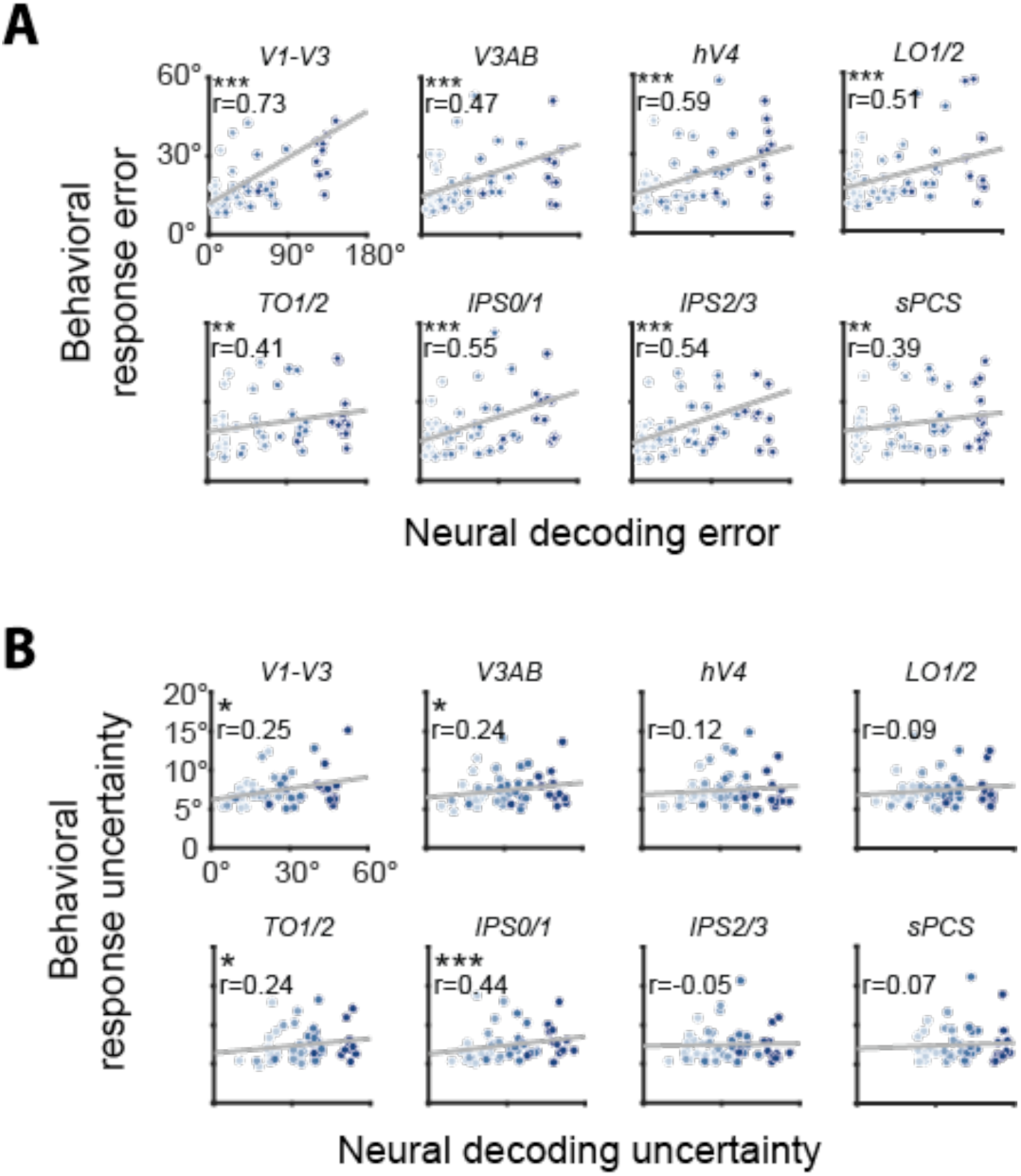
Brain-behavior correspondence. **(A)** Correlation between neural decoding and behavioral response The dots from lighter to darker indicate individual response sorted and binned according to individual neural decoding (see **Brain-behavior correspondence** for details). The gray lines the linear fits of the dots The stars in subplots indicate significant correlation between the neural decoding rror and behavioral response in all ROIs (all correlations ≥ .39; ps ≤ .003 Cohen’s ds ≥ 2.900). **(B)** Correlation between neural decoding uncertainty and behavioral response uncertainty. The dots from lighter to darker indicate individual response uncertainty sorted and binned according to individual neural decoding uncertainty (see **Brain-behavior correspondence** for details) The stars in subplots indicate significant correlation between the neural decoding uncertainty and behavioral response uncertainty in V1-V3 (r = .25, p .050 Cohen’s d = 18.759), V3AB (r = .24; p = .047, Cohen’s d = 2.535), TO1/2 (r = .24; p .048 Cohen’s d = .979), and IPS0/1 (r = .44; p < .001, Cohen’s d = 1.967)

**Figure S2.**
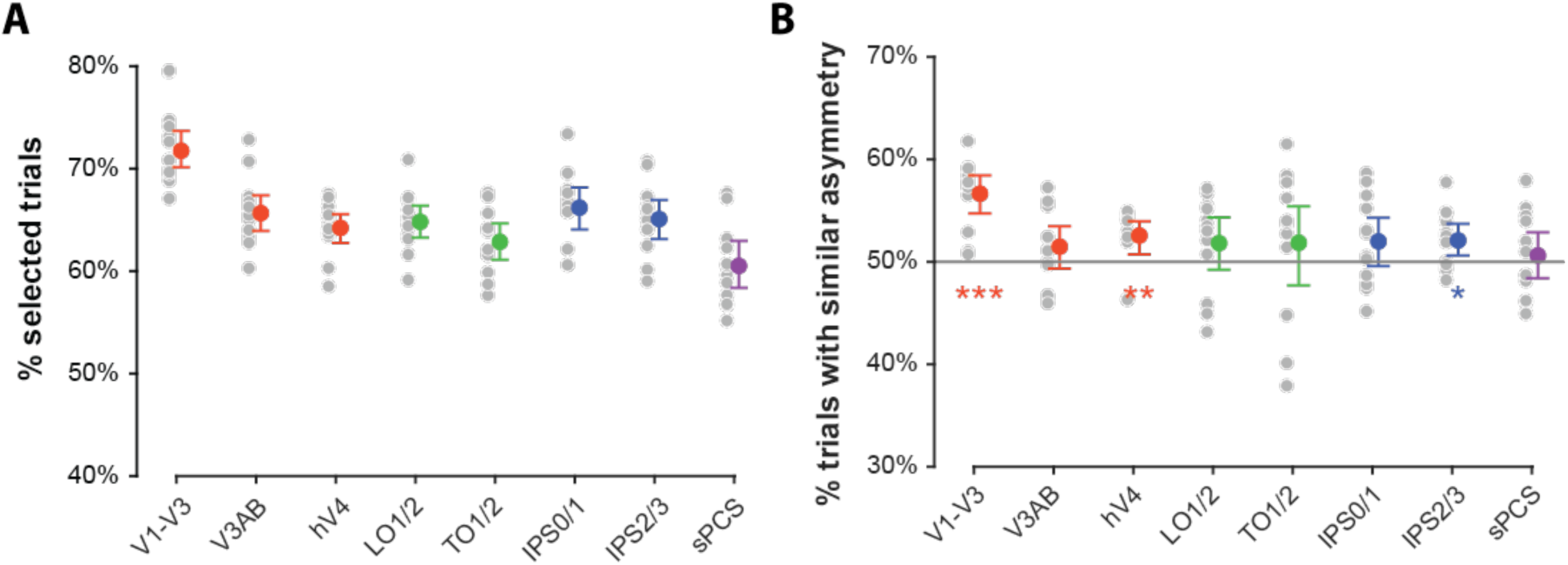
Brain-behavior correspondence restricted to trials with low distance between neural and behavioral We sought to rule out the possibility that the observed correspondence in distribution asymmetries (Figure 5) was driven by trials where the neural mea wa poor predictor of behavioral responses Noise could theoretically shift the neural away from the target and preserve higher likelihoods the target while behavioral responses remain clustered around the target We examined brain-behavior correspondence in Figure 5, limiting analysis to trials where the of the neural distribution within 1 SD of the of the bet distribution **(A)** Percentage of trials where the neural mean was within ne SD of the behavioral **(B)** Brain-behavior correspondence in these trials The correspondence quantified in Figure 5B Stars indicate significant correspondence in asymmetries in hV4 (p .009, Cohen’s d = .798), V1-V3 (p .001, Cohen’s d 1.937) and IPS2/3 (p = .018, Cohen’s d = .699).

**Figure S3.**
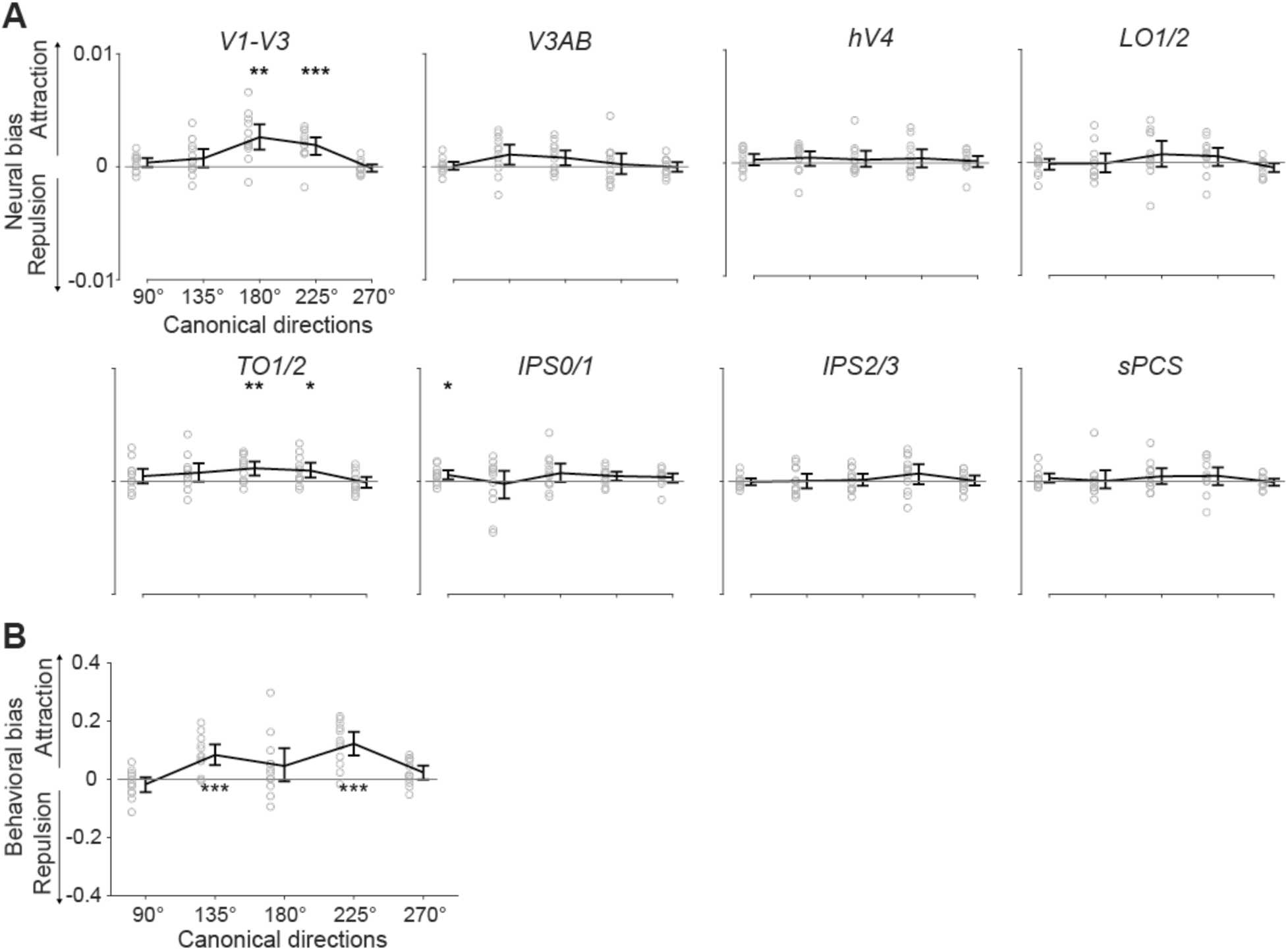
Asymmetry caused by categorial bias Trials grouped depending which canonical direction closest to the cued motion direction Each canonical direction labeled in polar angle space **(A)** Neural bias, quantified by calculating the likelihood over the motion direction of the nearby canonical direction and subtracting the likelihood the position in the neural distribution reflected about its Stars indicate significant attractive biases towards canonical directions in V1-V3 TO1/2, and IPS0/1 (B) Behavioral bias, quantified by subtracting the cumulative probability at the side towards the nearby canonical direction from the side away from the nearby canonical direction, relative to the first response Stars indicate significant attractive biases towards canonical directions in behavioral distributions

**Table S1.** Circular correlation between the decoded and target motion direction. For each subject and ROI, we report the uncorrected p-value obtained by comparing the actual circular correlation with null distribution by a permutation procedure (see Methods). A p-value less than 0.05 (highlighted in red) indicates significant circular correlation between decoded and target motion direction.

|  | V1-V3 | V3AB | hV4 | LO1/2 | TO1/2 | IPS0/1 | IPS2/3 | sPCS |
| --- | --- | --- | --- | --- | --- | --- | --- | --- |
| S1 | 0.43<br>(p=0.000) | 0.23<br>(p=0.000) | 0.12<br>(p=0.046) | 0.11<br>(p=0.060) | 0.10<br>(p=0.051) | 0.49<br>(p=0.000) | 0.39<br>(p=0.000) | 0.09<br>(p=0.069) |
| S2 | 0.26<br>(p=0.000) | 0.11<br>(p=0.040) | 0.09<br>(p=0.068) | 0.01<br>(p=0.449) | 0.08<br>(p=0.097) | 0.15<br>(p=0.005) | 0.21<br>(p=0.001) | -0.02<br>(p=0.638) |
| S3 | 0.40<br>(p=0.000) | 0.24<br>(p=0.001) | 0.15<br>(p=0.019) | 0.12<br>(p=0.074) | -0.12<br>(p=0.943) | 0.29<br>(p=0.000) | 0.31<br>(p=0.000) | -0.06<br>(p=0.782) |
| S4 | 0.51<br>(p=0.000) | 0.24<br>(p=0.000) | 0.20<br>(p=0.001) | 0.29<br>(p=0.000) | 0.24<br>(p=0.000) | 0.24<br>(p=0.000) | 0.19<br>(p=0.001) | 0.05<br>(p=0.219) |
| S5 | 0.41<br>(p=0.000) | 0.18<br>(p=0.000) | 0.12<br>(p=0.042) | 0.23<br>(p=0.000) | 0.18<br>(p=0.000) | 0.20<br>(p=0.001) | 0.32<br>(p=0.000) | 0.16<br>(p=0.008) |
| S6 | 0.28<br>(p=0.000) | 0.05<br>(p=0.264) | 0.01<br>(p=0.496) | 0.03<br>(p=0.356) | 0.01<br>(p=0.451) | -0.10<br>(p=0.901) | 0.12<br>(p=0.054) | 0.12<br>(p=0.065) |
| S7 | 0.17<br>(p=0.008) | -0.01<br>(p=0.557) | 0.05<br>(p=0.218) | 0.05<br>(p=0.218) | 0.11<br>(p=0.048) | 0.00<br>(p=0.478) | 0.11<br>(p=0.051) | 0.07<br>(p=0.135) |
| S8 | 0.46<br>(p=0.000) | 0.44<br>(p=0.000) | 0.19<br>(p=0.000) | 0.35<br>(p=0.000) | 0.21<br>(p=0.000) | 0.23<br>(p=0.000) | 0.28<br>(p=0.000) | 0.07<br>(p=0.138) |
| S9 | 0.36<br>(p=0.000) | 0.11<br>(p=0.047) | 0.18<br>(p=0.001) | 0.38<br>(p=0.000) | 0.03<br>(p=0.296) | 0.08<br>(p=0.097) | 0.18<br>(p=0.003) | -0.08<br>(p=0.893) |
| S10 | 0.38<br>(p=0.000) | 0.22<br>(p=0.000) | 0.11<br>(p=0.051) | 0.17<br>(p=0.002) | 0.03<br>(p=0.352) | 0.18<br>(p=0.001) | 0.10<br>(p=0.071) | -0.06<br>(p=0.821) |
| S11 | 0.34<br>(p=0.000) | 0.19<br>(p=0.002) | 0.08<br>(p=0.119) | 0.09<br>(p=0.101) | -0.05<br>(p=0.780) | 0.21<br>(p=0.003) | -0.06<br>(p=0.834) | -0.01<br>(p=0.550) |
| S12 | 0.79<br>(p=0.000) | 0.81<br>(p=0.000) | 0.73<br>(p=0.000) | 0.73<br>(p=0.000) | 0.56<br>(p=0.000) | 0.62<br>(p=0.000) | 0.65<br>(p=0.000) | 0.25<br>(p=0.000) |

